# hnRNPA1 RGG-box acts a ”catalyst” in UP1-induced destabilization of RNA and DNA telomeric G-Quadruplexes

**DOI:** 10.1101/2023.08.21.554222

**Authors:** Sangeetha Balasubramanian, Irawati Roy, Rajeswari Appadurai, Anand Srivastava

**Affiliations:** Molecular Biophysics Unit, Indian Institute of Science, Bangalore, Karnataka 560012, India; Department of Biology, Indian Institute of Science Education and Research, Tirupati, Andhra Pradesh 517619, India

**Keywords:** hnRNPA1-LCD, RNA and DNA G-quadruplex, telomere maintenance, inte-grative modeling, multidomain IDPs

## Abstract

hnRNPA1, a protein from the heterogeneous-nuclear ribonucleoprotein family, mediates cellular processes such as RNA metabolism and DNA telomere maintenance. Besides the folded RNA recognition motifs, hnRNPA1 has a ∼ 135 amino-acids long low-complexity domain (LCD) consisting of an RGG-rich region and a prion-like domain (PrLD). Biochemical data suggest that RGG-rich region modulates recognition of G-quadruplexes (GQs) in the telomeric repeats. Here, we utilize an in-house developed replica exchange technique (REHT) to generate the heterogeneous conformation ensemble of hnRNPA1-RGG and explore its functional significance in telomere maintenance. Single chain statistics and abundance of structural motifs, as well as consistency with experimentally reported structural data, suggest faithful recapitulation of local interactions. We also introduce a protocol to generate functionally significant IDP-nucleic acid complex structures that corroborate well with the experimental knowledge of their binding. We find that RGG-box preferentially binds to the grooves and loops of GQs providing specificity towards certain GQ structures with its Phe, Tyr, and Asn residues forming essential hydrogen bonds and electrostatic interactions. Several of these residues were also identified as important by the reported HSQC chemical shift data. Our binding and simulations studies also revealed that a minor population of the RGG-box can destabilize telomeric GQs, thereby expediting the unfolding activities of hnRNPA1-UP1 at the telomeric end.

## I. INTRODUCTION

Heterogeneous nuclear ribonucleoprotein A1 (hnRNPA1) is a member of a complex and diverse group of proteins called hnRNPs that collectively play an important role in processing heterogeneous nuclear RNAs into mature mRNAs. This abundant nuclear protein modulates a number of RNA metabolism processes, including transcription^1–3^, splicing^3–6^, RNA stability, export^7–9^ and translation^10–12^. However, the role of hnRNPA1 is not just limited to mRNA bio-genesis but extends to other processes such as microRNA processing and telomere maintenance^13^. Structurally (Fig. 1(a)), the N-terminus of the protein contains two well-folded RNA recognition motifs (RRMs) connected by a small linker region. Jointly these two motifs constitute Unwinding protein 1 (UP1)^14^, which mediate interac-tions with target RNAs. UP1 domain is shown to bind to and stablize telomeric ssDNA and also unfold telomeric G-quadruplexes (GQ)^13,15–18^. In addition, a flexible Glycine rich C-terminal region consisting of Arg-Gly-Gly tripeptide repeats (RGG), interspersed with aromatic (Phe, Tyr) residues to form RGG boxes, provide both protein and RNA binding capabilities to hnRNPA1^14^. The RGG-box domain of hnRNPA1 is a low-complexity domain with limited amino acid composition (Fig. 1 b). Downstream from the RGG boxes, the C-terminal domain harbours a prion-like domain (PrLD) and a nuclear shuttling sequence called the M9 sequence^19^. PrLD has been shown to induce protein-protein interaction, leading to stress granule assembly and pathological protein aggregation^19^. The RGG and PrLD domain together constitute the disordered low complexity domain (LCD) of hnRNPA1.

**FIG. 1:**
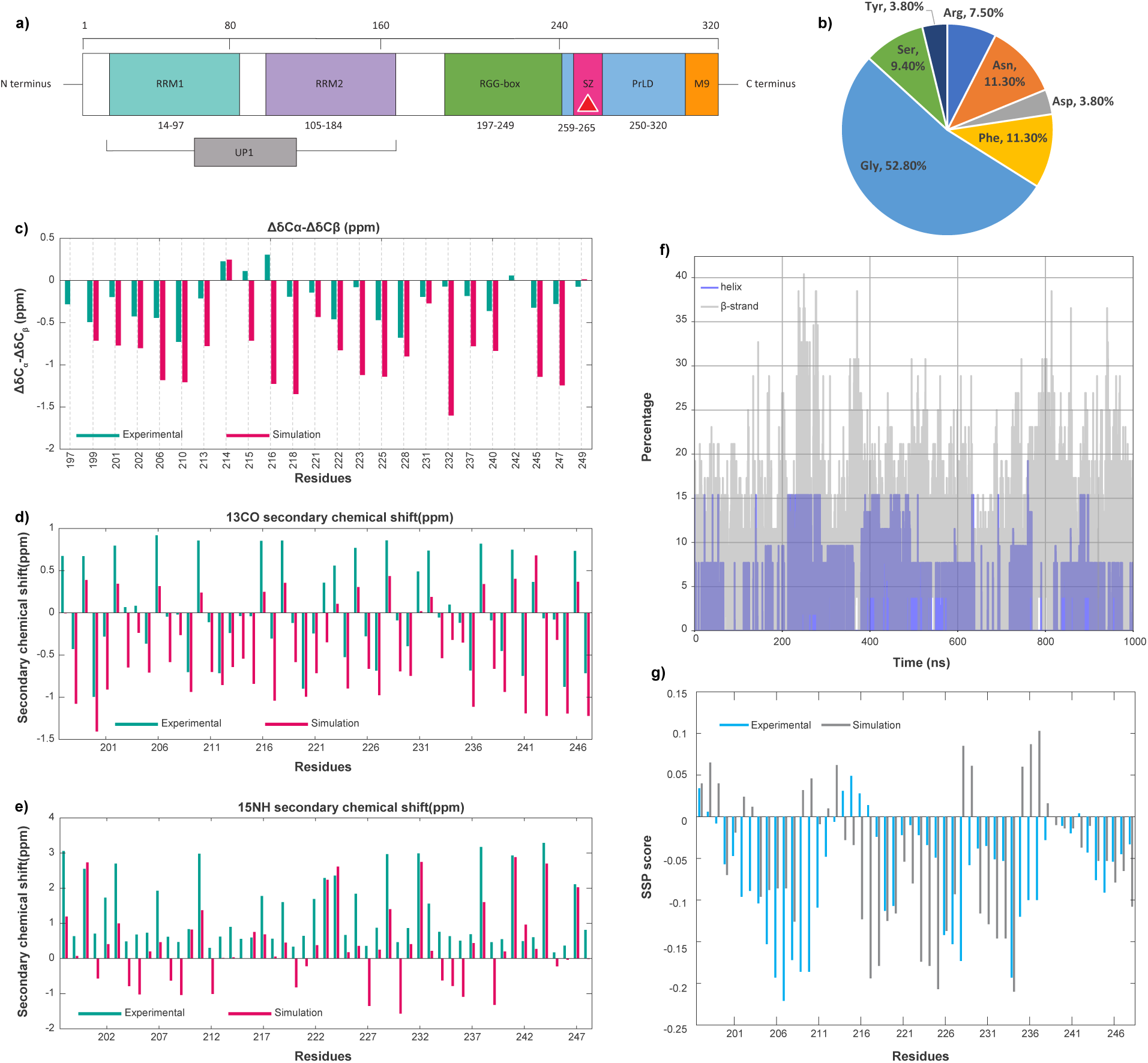
a) Domain architecture of hnRNPA1. The Teal and Purple coloured domains are the N terminal ordered RNA recognition motifs. Together, these two domains are called UP1. The steric-zipper motif (259-265) is coloured in Magenta. The Green and the Blue domains are intrinsically disordered RGG-box domain and PrLD respectively. The Orange coloured domain is a nuclear shuttling sequence called M9. b) Sequence compostion of RGG-box motif. c), d), e) are the *δ*C*α*-*δ*C*β*, ^13^*CO* and ^15^*N ^H^* secondary chemical shift plots for RGG, respectively. Teal is experimental data. Dark-pink indicates simulation data. The *δ*C*α*-*δ*C*β* chemical shifts are calculated and plotted for the residues containing *Cβ* atoms. f) Comparison of *α*-helix (grey) and *β*-sheet (blue) content of RGG during the simulation. g) Predicted secondary structure propensity scores from chemical shift data of experiments (cyan) and simulation (grey) using the SSP algorithm. SSP score above 0 indicates helical propensity. SSP score below 0 indicates *β*-sheet propensity.

Several lines of evidence have demonstrated that both UP1 and RGG-box domains are involved in GQ unfolding activity at the telomeric end of DNA^20,21^. It is believed that the RGG-box interacts with the loop nucleotides of the GQ structure, facilitating the initial specific recognition and binding of UP1 to the GQ DNA^20^. As such, RGG-box acts as an adaptor for the UP1 by bringing it closer to the GQ, which leads to an efficient binding of UP1 followed by unfolding of the GQ structure^20^. Finally, UP1 forms a high-affinity complex with the single-stranded DNA and stabilises the extended form. Besides DNA, very recent studies have also reported structure-dependent recognition of RNA GQ by the RGG-box of hnRNPA1, where the long non-coding RNA called the telomeric repeats containing RNA (also known as telomere RNA or TERRA) forms GQ structures and has regulatory functions over telomere replication. It was shown that the RGG-box of hnRNPA1 binds to the loops in TERRA GQ but not with the single-stranded RNA, indicating that hnRNP can specifically recognize a GQ structural motif^22^. Despite tremendous strides made in the mechanistic understanding of hnRNPA1 interactions with GQ using biochemical studies, high-resolution structural and dynamic insights into hnRNPA1-GQ interaction and telomere maintenance are still missing. In our work, we utilize the integrative molecular modeling approach that combines atomic-resolution molecular simulations with available ensemble averaged data from NMR to faithfully reconstruct the atomic-resolution ensemble of hnRNPA1-RGG and use the structural data to investigate the functional role of RGG in telomere maintenance.

In this work we have focused on and explored the structural heterogeneity and associated biological activities of hnRNPA1 with the help of our advanced-sampling^23^ and large scale biomolecular simulations of hnRNPA1-RGG and its RNA/DNA GQ complexes. The paper is organized as follows. In the Results and Discussion section following this Introduction section, we report the data from our extensive docking and simulations of RGG-region interacting with multiple RNA and DNA GQ topologies. Our simulations provide molecular-level insights into the reported role of RGG-region in telomere maintenance. We discuss in detail how the RGG-region binds differently to telomeric DNA and telomeric RNA GQs and we also demonstrate how a very small population of hnRNPA1-RGG conformers are capable of destabilizing the GQs. After the Results section, we briefly summarize our observations and findings in the Conclusions section, which is followed by the Materials and Methods where we provide details of all the methods that we have used. Additional information is included in the Supporting Information (SI) file. All our data including the simulated trajectories and input files for our simulations, all analyses related data and codes are publicly available here.

## II. RESULTS AND DISCUSSION

### A. Atomic resolution structural insights into the RGG conformation ensemble

#### 1. Conformations consistent with NMR data show prevalence of β-motifs

We obtained a heterogeneous ensemble of the RGG domain (197-249 amino acids) (1(a)) to gain conformational insights by harnessing a Hamiltonian and temperature-scaled hybrid replica exchange technique called Replica Exchange with Hybrid Tempering (REHT)^23–25^. Please refer to the Methods section for more details. We calculated the ΔC*α*-ΔC*β*, ^13^*CO* and ^15^*N ^H^* secondary chemical shifts using the conformation ensemble generated from our REHT simulation. The residue-wise comparison between the calculated data from the simulation and the experimentally-available chemical shift data showed a considerable match^26^ as shown in Fig. 1(c,d,e). Upon performing a basic secondary structure analysis across the time evolution of our REHT ensemble, we observed a significant propensity to form transient secondary structures, preferentially *β*-sheet and extended *β*-structures at different regions of RGG, as shown in Fig S1 in SI. Moreover, the overall percentage of *β*-structures in each frame is higher when compared to *α*-helices (Grey in Fig. 1(f)). To verify this, we also used the chemical shift data of the experiment and simulation ensemble to predict the secondary structure propensities per residue. The secondary structure prediction (SSP)^27^ scores of both experimental and simulated data re-emphasise our secondary structure findings of the ensemble data by showing a higher *β*-sheet population compared to that of *α*-helix (Fig. 1(g)). Altogether, the secondary chemical shifts and secondary structure analysis suggest a heterogeneous ensemble with an overall inclination to form local contacts through transient secondary structure formation.

At this point, it may be pertinent to discuss the prevalent ambiguity regarding secondary structure prediction from chemical shift data using conformational ensemble generated from molecular simulations. First of all, at the experimental interpretation level itself, predicting the secondary structure of intrinsically disordered proteins (IDPs) from chemical shift data is often potentially ambiguous due to several key reasons. Some of these reasons are the lack of a well-defined secondary or tertiary structure under the physiological conditions of IDPs and IDRS, the chemical shift averaging over many conformations, the high dependence of the chemical shift of an IDP on its local environment, lack of suitable reference database and last but not the least, the weak secondary structure propensities. The weak and transient secondary structure propensities exhibited by IDPs often do not produce chemical shift deviations that are strong enough to be reliably detected and interpreted using conventional secondary structure prediction algorithms. Due to this specific reason, in our particular case of RGG, we have observed outcome from SSP analysis that has to be interpreted in the light of inherent ambiguity in making strong deductions from the experimental data itself. Use of ^13^*C_α_*, ^1^3*C_β_* and ^13^*CO* gave a higher percentage of *α*-helix propensity, whereas use of ^13^*C_α_*, ^1^3*C_β_*and ^1^*H_α_* produced a higher *β*-sheet propensity. Due to the effect of the weak secondary structure propensity, the reliability of the prediction gives way to obvious ambiguity. Nevertheless, in this work, we followed the recommended set of raw chemical shift data according to the SSP prescription of IDP (which are ^13^*C_α_*, ^1^3*C_β_* and ^1^*H_α_*) and were able to qualitatively match the result with the outcome of secondary structure calculation from our simulation data set using DSSP.

### B. Functional insights from RGG conformation ensemble: Role of RGG box in telomere maintenance

In 2018 Ghosh and Singh compared the 15N-1H HSQC data of the UP1+RGG-box domain and the isolated RGG-box domain (^20^). A superposition of the NMR spectra of these two constructs showed that RGG occupies the same conformational space with or without the existence of UP1. This in turn sheds light on the modular nature of the UP1-RGG construct where the tethered RGG-box behaves as an structurally independent domain from the folded UP1 domain. This also opens up the possibility that the full-length LCD (RGG+PrLD) could be modular. Bionformatics studies on sequence space has suggested distinct compositional boundaries within IDRs – revealing a possible non-random modular organization in the sequence space^28^. Fortuitously, NMR studies from completely different groups have reported the HSQC spectra of the RGG domain^20^ and the full-length LCD with the RGG domain and PrLD domain together as one chain^26^. In order to explore the possibility that full-length LCD of the hnRNPA1 could be modular, we superposed these HSQC spectra and it is clearly seen that the chemical shifts of RGG overlaps in the two systems (Figure S2 (a) in SI). The NMR data clearly portrays the modularity of hnRNPA1 and signifies that the RGG domain, in spite of being disordered, can function independent of the other regions of hnRNPA1 such as UP1 and PrLD. This structural and functional modularity motivated us to dig deeper into the functionality of the RGG domain, in particular, our focus was to understand the functional relevance of RGG box domain in telomere maintenance. hnRNPA1 expresses a concentration-dependent binding with telomeric RNA and DNA; in particular, it localizes at the GQ structural motif. Studies have hypothesised that the RGG-box binds to GQ, recruits UP1 which in turn destabilises the GQ^20,22^. We explore this hypothesis of RGG-GQ interactions, but the heterogeneity of the RGG ensemble complicates the selection of a reference RGG structure for the study. We used our recently developed machine learning-based clustering technique, which employs t-distributed stochastic neighbour embedding (t-SNE), to group the huge heterogeneous ensemble into 50 homogeneous clusters^29^ (see Fig. S2(b) in SI). Our tSNE method clustering algorithm for IDPs combines t-distributed stochastic neighbor embedding projection and kMeans clustering technique to group the heterogeneous data into homogenous subgroups. Through our Silhouette score, we can see that each cluster is evidently highly homogeneous, whereas the conformations between clusters are heterogeneous. The algorithm and parameters used to perform tSNE clustering is discussed in the Methods section. We selected a central conformation from each cluster to represent the different states sampled by the RGG in solution and further used them to study the functional interactions. Being an IDP, RGG might interact with other partners via several mechanisms like “folding upon binding” or fuzzy interactions^30–32^. In computational studies, first-pass docking methods generally provide insight into the formation of “encounter complexes”. To identify these encounter complexes and the probable basis for GQ recognition by RGG, we performed docking studies of RGG with telomeric DNA as well as RNA quadruplexes using HADDOCK. Further, the docked structures were then subjected to molecular simulations to capture the dynamics of the complex. A flowchart depicting the protocol adopted to generate the initial RGG-GQ encounter complexes is shown in Fig. 2.

**FIG. 2:**
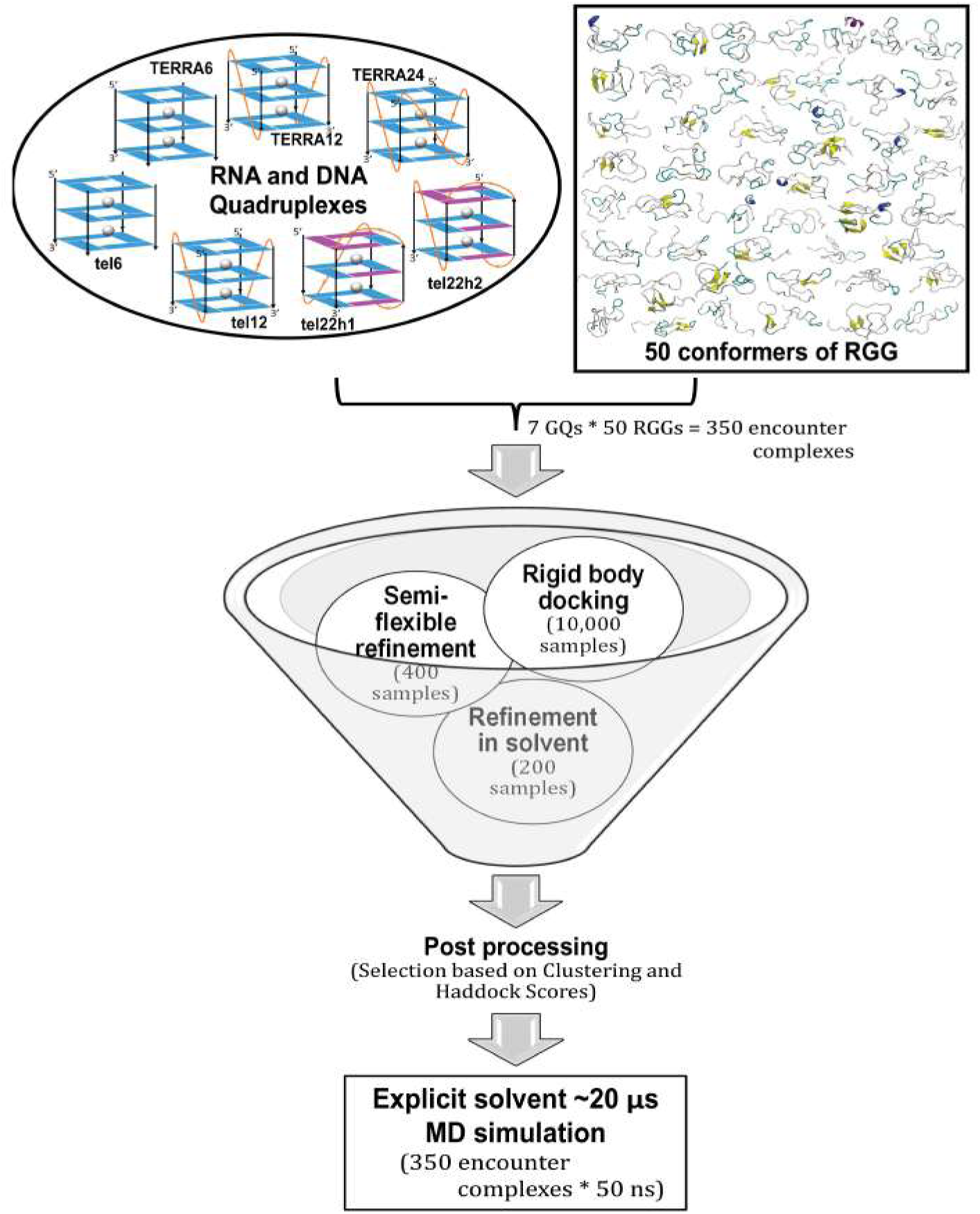
Flowchart of the protocol adopted to generate RGG-GQ complexes. Seven different telomeric DNA and RNA GQ structures chosen from literature were docked with each of the 50 conformers of RGG obtained by t-SNE clustering of the generated ensemble. HADDOCK protocol with center of mass distance criteria was used to reduce the sample space from 10,000 initial random geometries for each complex. The 350 (7 GQs with 50 RGGs) encounter complexes were further simulated for 50 ns each resulting in 17.5 *µ*s ensemble to explore the mode of binding, stability and conformational dynamics of RGG-GQ binding process.

#### 1. Encounter complexes of telomeric RNA GQ are ”end stacking” and telomeric DNA GQ are “groove binding”

We docked each of the 50 conformations of RGGs with 7 different unimolecular, bimolecular and tetramolecular GQs of telomeric RNA (named TERRA6, TERRA12 and TERRA24, respectively) and DNA (named tel6, tel12, tel22h1 and tel22h2). The systems under consideration are listed in Table I, and the HADDOCK scores for all complexes are plotted in Fig. S3 in SI. Preliminary visual inspection and analysis of three-dimensional occupancy maps (drawn on an isosurface with a 6 °A distance cutoff) reveal the most preferred binding sites on the 7GQs by the 50 RGG conformations (Fig S3 in SI). The maps also clearly distinguish the mode of binding of RGG with RNA and DNA GQs. The RGG dominantly encounters the solvent-exposed external G-quartets of TERRA RNA in an ”end stacking” mode. In contrast, it binds mostly in the grooves and loops of DNA GQs.

**TABLE I:** Telomeric RNA and DNA structures modeled in this study. For every GQ, there are fifty complexes.

| Systems | PDB ID | Apo (ns) | RGG-GQ(ns) | Comments |
| --- | --- | --- | --- | --- |
| **TERRA6** | based on 6GE1 | 300 | 50 | tetramolecular RNA |
| **TERRA12** | 2KBP | 300 | 50 | bimolecular RNA |
| **TERRA24** | based on 1KF1 | 300 | 50 | unimolecular RNA |
| **tel6** | 1NB9 | 300 | 50 | tetramolecular DNA |
| **tel12** | 1K8P | 300 | 50 | bimolecular DNA |
| **tel22h1** | 2HY9 | 300 | 50 | unimolecular DNA hybrid 1 |
| **tel22h2** | 2JPZ | 300 | 50 | unimolecular DNA hybrid 2 |

Interestingly, the binding modes influence the relative binding strength of the complexes. The calculated HADDOCK scores for the complexes in end stacking mode, such as those involving TERRA6, TERRA12 and TERRA24, are relatively lower in the range of -85 to -65. Whereas in groove binding mode, as in tel6, tel12, and tel22 complexes, the binding is slightly better (in the range of -100 to -75), with tel12 showing the strongest binding owing to the mixed mode of interactions. Another significant observation is that the HADDOCK scores for complexes involving tetra-molecular GQs (TERRA6, tel6) without propeller loops are weaker than those for GQs with propeller loops (such as TERRA 12, TERRA24, tel12 and tel22). These results corroborate well with experimental findings that the RGG primarily interacts with the GQ loops^20,22^. Our atomic resolution complex structures further provide deeper insights into the residue-wise contributions to the inter-molecular binding. In Tables S1 and S2 we provide a detailed list of inter-molecular interactions for the three strongest RGG-GQ complexes, screened by their HADDOCK scores. Intriguingly, in all these complexes, we observe overwhelming contributions of ARG and GLY from the RGG repeat motifs to the Hydrogen bonding and electrostatic interactions, emphasizing the importance of RGG repeats in nucleic acid recognitions. Also the interspersing aromatic residues such as TYR237, TYR244 and several PHE residues are mediating the *π*-stacking interactions. We also note a larger contribution of SER residues in the DNA complexes.

#### 2. Both DNA and RNA GQ with loops and overhangs have stronger binding consistent with recent biochemical observations

All the encounter complexes obtained from HADDOCK were simulated for a period of 50 ns each (a total of 17.5 *µ*s of all-atom simulations) to monitor the variations in RGG binding orientation and its induced changes in the GQ structural stability. These simulation trajectories clearly revealed the dynamic changes in the binding modes of RGG with the GQs and the type of stabilizing interactions. Fig. 3 depicts the inter-molecular interactions between the amino acids in RGG and each base of GQ. These interactions are normalized with the individual amino acid composition and indicate the participation of each amino acid type towards the stability of the complex. When an amino acid in RGG resides within 4 °Aof the central Guanine O6 atom, it is considered to explore the end-stacking mode. Groove binding mode is identified when an amino acid of RGG forms contact with any of the atoms exposed in the grooves, like ”N2”, ”C2”, ”N3”, ”C4”, ”N6”, ”C8” or ”N9” atoms of Guanine. Similarly, the loop binding mode of RGG is described by the interaction with TTA, irrespective of its backbone, sugar or base. The three binding modes are not exclusive, and a given conformation of RGG might exhibit all three binding modes simultaneously by different regions of RGG. The encounter complexes from docking studies clearly show end stacking mode of TERRA12, TERRA24 and tel12, and groove or loop binding modes in other GQ complexes. However, after simulations, the binding modes vary and groove or loop binding modes are preferred in all GQ complexes Fig. 4.

**FIG. 3:**
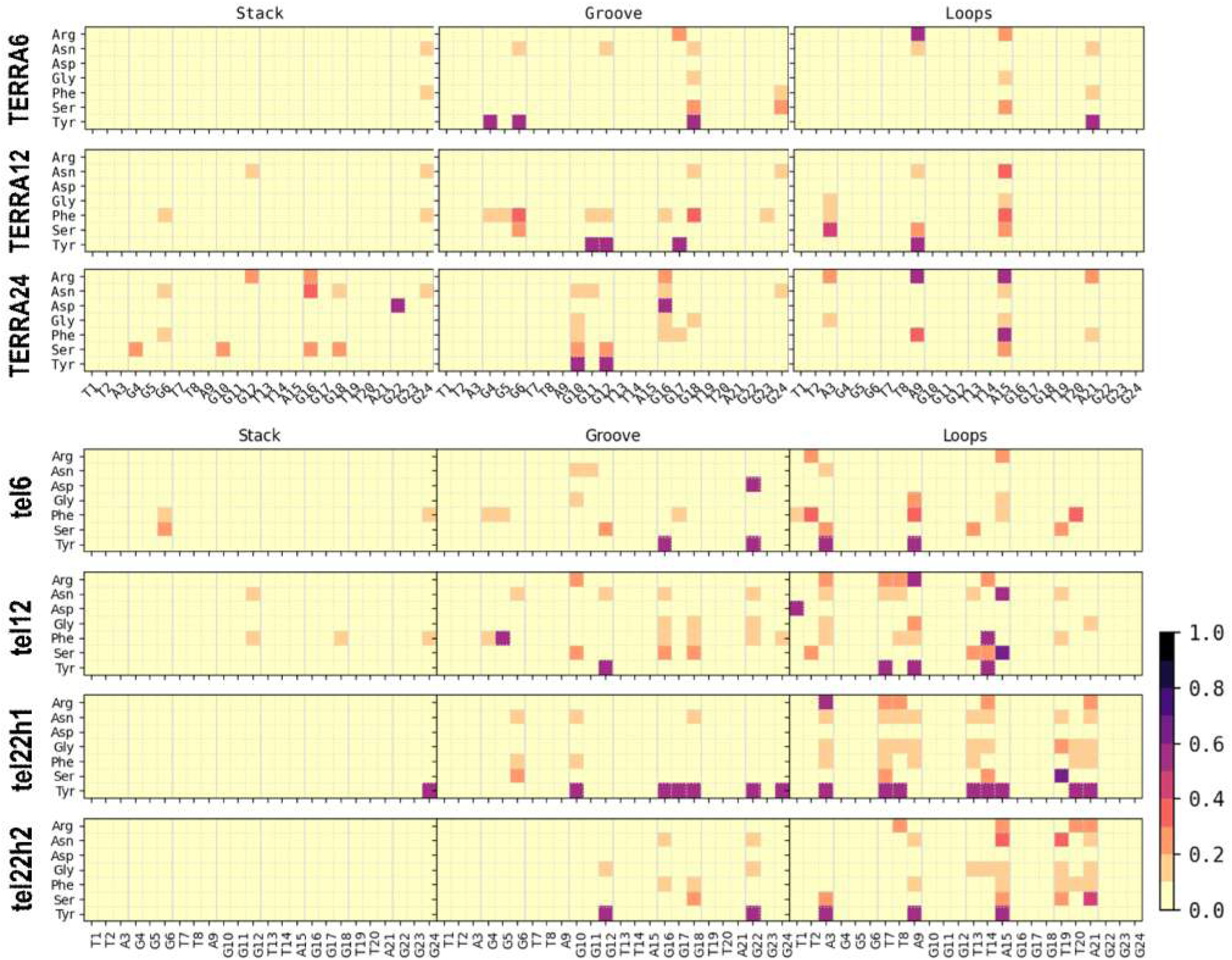
Binding modes of RGG with GQ. The normalized number of interactions between each amino acid type in RGG (y-axis) with every base of GQ (x-axis) was calculated from the simulated complexes. The interactions were calculated based on the distance between heavy atoms and the number of interactions were normalized on the total number of each amino acid type. Rows represent the seven GQ systems, while the columns represent end-stacking, groove and loop binding modes. The contact probabilities are color coded on a yellow to black scale where darker colors indicate a higher probability of interactions.

**FIG. 4:**
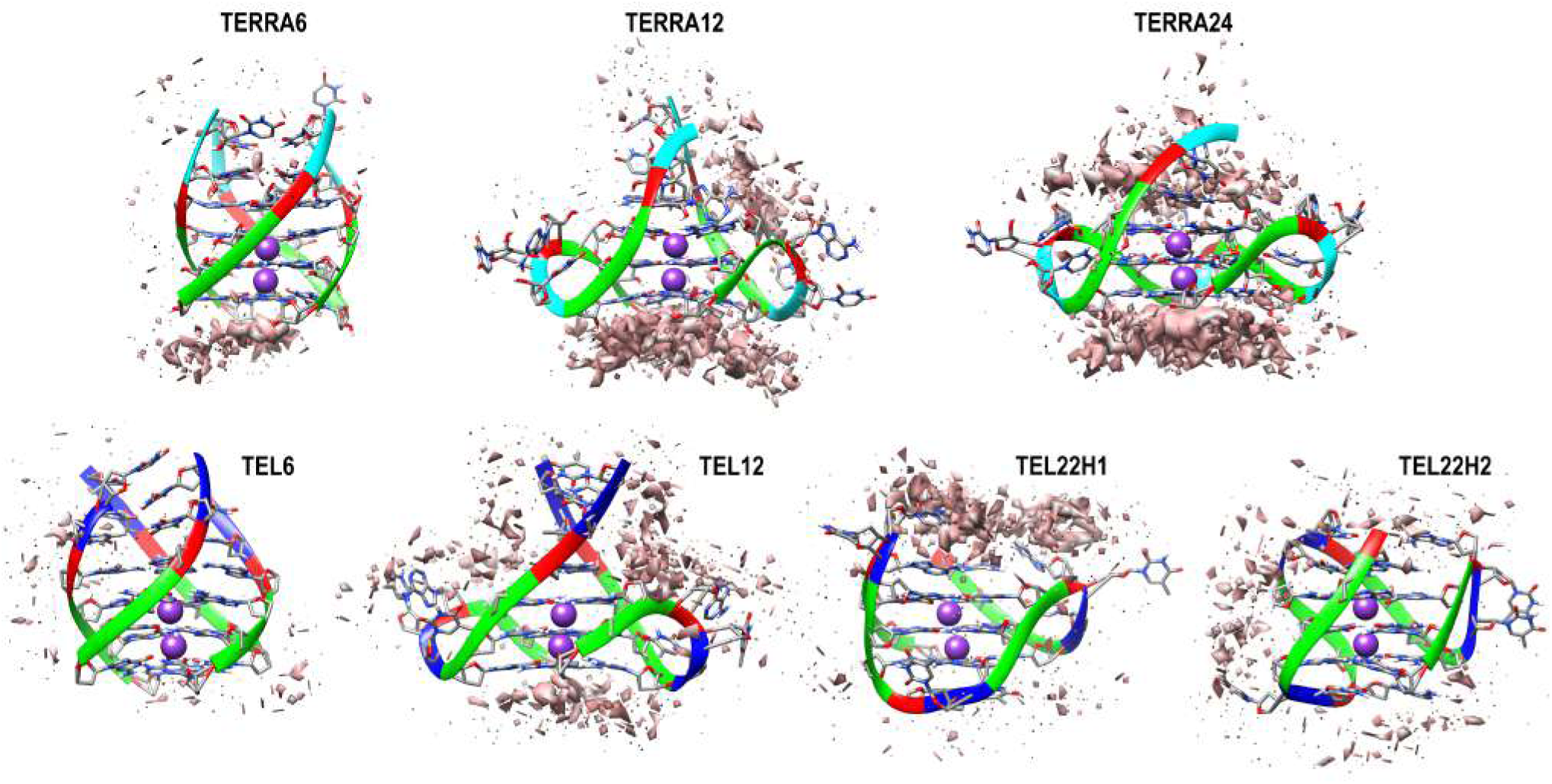
Occupancy map depicting the extent of RGG-GQ interaction after the 50ns simulation. High density of interactions at specific sites on GQ are shown as an isosurface mimicking the first solvation shell (orange surface). The interacting sites and density varies considerably from the complexes before simulation (see Fig S3 in SI for occupancy maps before simulation). The GQ backbone is shown as ribbons while the bases are displayed as sticks. The bases are colored as Green: Guanine, Red: Adenine, Blue: Thymine and Cyan: Uracil.

Fig. 3 clearly shows that in the telomeric DNA GQ complexes, interactions with grooves and loops are equally predominant while in case of telomeric RNA GQ complexes, the in-teractions with grooves out-number those with the loops. Interestingly, groove and loop binding modes dominate in TERRA24 and tel12 complexes after simulation, unlike the stacking interactions in encounter complexes. Among the RGG residues, Arg and Asn prefer to interact with the TTA loops of telomeric DNA GQs. However, Phe and Tyr contribute towards interactions with both Guanines and the TTA loops.

#### 3. Simulated ensemble of RGG-GQ complexes show interactions consistent with NMR experiments

In the previous section, we delineated the RGG binding sites in GQ and the preference of each amino acid type to interact with specific binding sites. Following this, we explore the lifetime of all RGG-GQ interactions by generating contact matrices (Fig. 5 and 6). The lifetime of RGG-GQ interactions in TERRA complexes is higher than DNA GQ complexes. Preliminary inspection of these contact maps clearly show that the N-and C-termini of RGG do not exhibit strong interactions. Previous NMR studies have highlighted the role of Tyr244 in binding GQs. Other residues of RGG like Ser197, Ser231, Arg232, Gly233, Tyr244, Asn245 and Gly246 were also reported to show chemical shift perturbations upon binding with GQ^20,22,26^. Our contact analysis shows that these residues interact strongly with GQs, along with several other residues, particularly in the core of RGG (between Phe216 to Arg232). Altogether, by using our protocol, we were able to generate protein-nucleic acid complex structures that match well with experimental data and also provide additional atomic-level insights into the structural features.

**FIG. 5:**
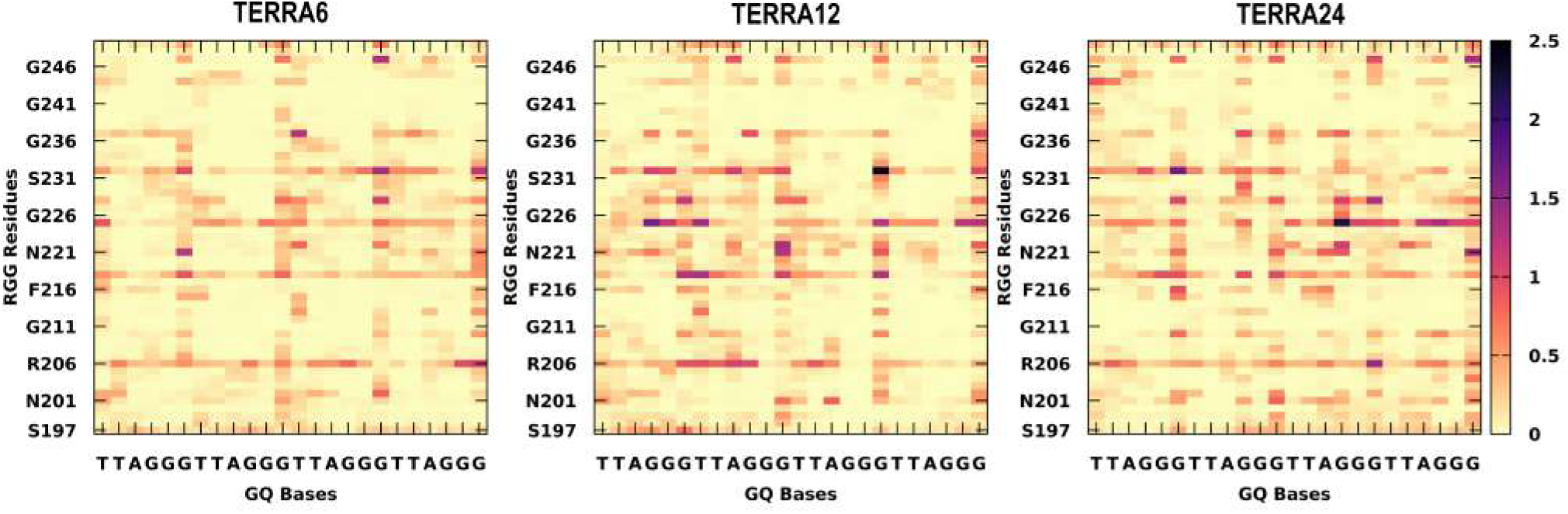
Residue-wise contact maps representing RNA-GQ interactions. Columns represent the three TERRA complexes and rows represent the 7 different amino acids in RGG. x-axis indicates the GQ bases while the entire RGG sequence is shown in y-axis. The color scale varies as white to dark blue to indicate longer lifetime of interactions. The contacts were calculated with a distance cutoff of 4 Å.

**FIG. 6:**
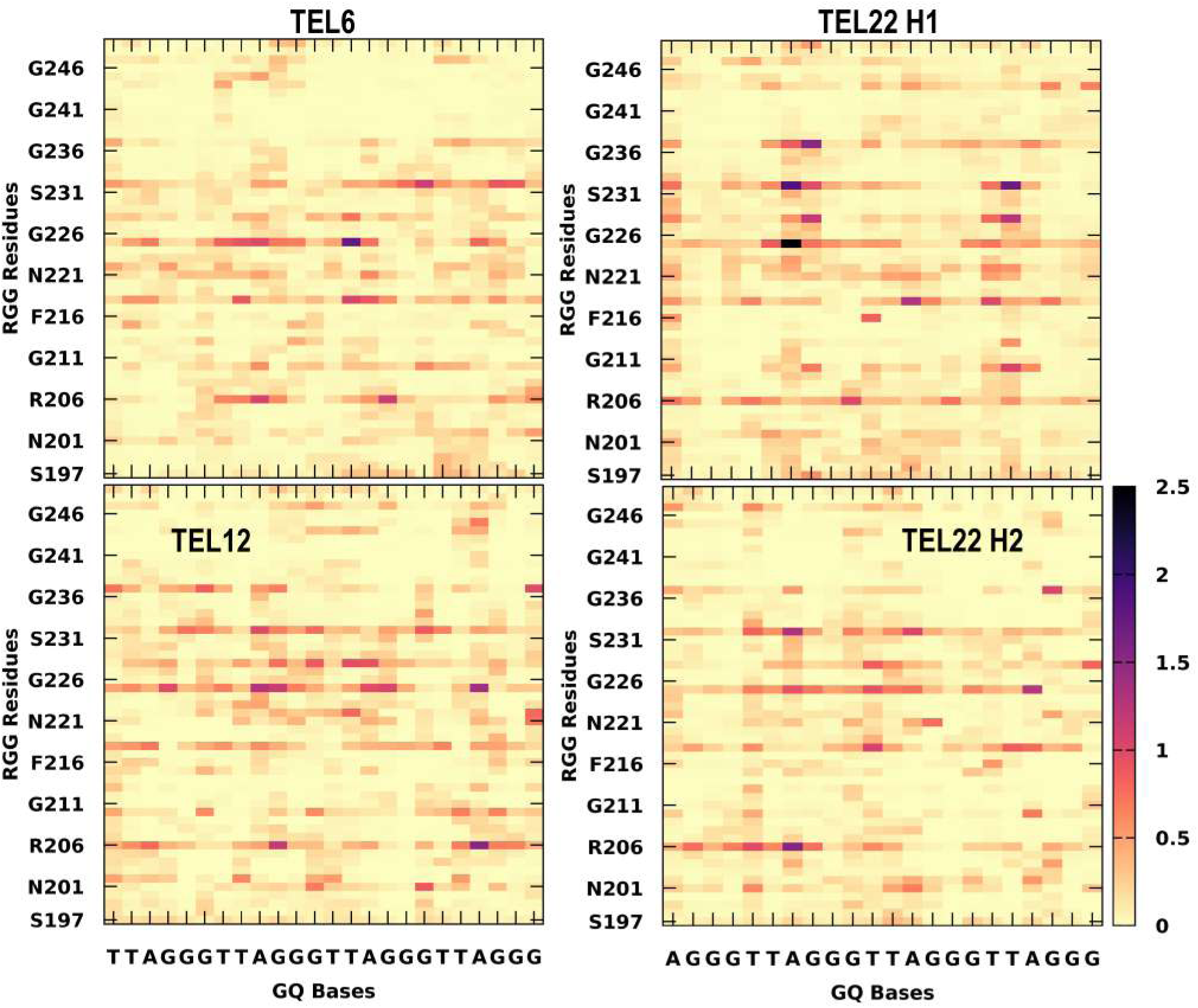
Residue-wise contact maps representing DNA-GQ interactions. Columns represent the four telomeric DNA GQ complexes and rows represent the 7 different amino acids in RGG. x-axis indicates the GQ bases while the entire RGG sequence is shown in y-axis. The color scale varies as white to dark blue to indicate longer lifetime of interactions. The contacts were calculated with a distance cutoff of 4Å.

#### 4. RGGs are able to destabilize both RNA and DNA GQs in a non conformation-specific manner

Among the 350 simulated complexes, visual inspection shows that few of the GQ complexes were unstable. Hence, before analyzing these unstable complexes, we ensured the stability of the GQs by simulating their unbound conformations. The human telomeric repeat sequence has three consecutive guanines (d(TTAGGG)*_n_*), and consequently, each GQ topology chosen in our study has three G-quartets. Each G-quartet is stabilized by 2 Hoogsteen H-bonds between adjacent guanines and a total of 8 H-bonds per quartet. Collectively, the three quartet GQs are stabilized by a minimum of 24 H-bonds. Our apo-state simulations show that all seven GQ structures were stable for the entire simulation period of 300 ns as seen by the time evolution of their G-quartet hydrogen bonds (Fig. S5 in SI). Hence, any changes in the stability of GQs in the RGG bound complexes are considered as the effect of RGG binding. During simulation, the RGG remains dynamic and experiences changes in its conformation as well as its binding with GQ. Considering the stability of apo GQ structures and the dynamic behavior of RGG, if a 50 ns simulation time induces any local disruptions in the GQ structure, it can be interpreted as a direct effect of RGG binding. The average number of hydrogen bonds between RGG and GQ as well as within the G-quartets were calculated in all the simulated complexes and a scatter plot correlating these is shown in Fig. 7. A majority of the GQs possess 22 ± 2 hydrogen bonds and hence, they are stable. Loss of even 2 hydrogen bonds in each of the three G-quartets would elicit destabilization in the overall GQ structure. Therefore, we used an arbitrary cut-off of 15 hydrogen bonds or less (of the total 24) within the G-quartets to consider it as unstable. Fig. 7 shows that at least one among each of the TERRA complexes is unstable. Similarly, in case of the telomeric DNA (Fig. 7), three tel6 and one tel22h2 complexes are unstable. Snapshots of these complex structures after the 50 ns molecular dynamics (MD) simulation are shown.

**FIG. 7:**
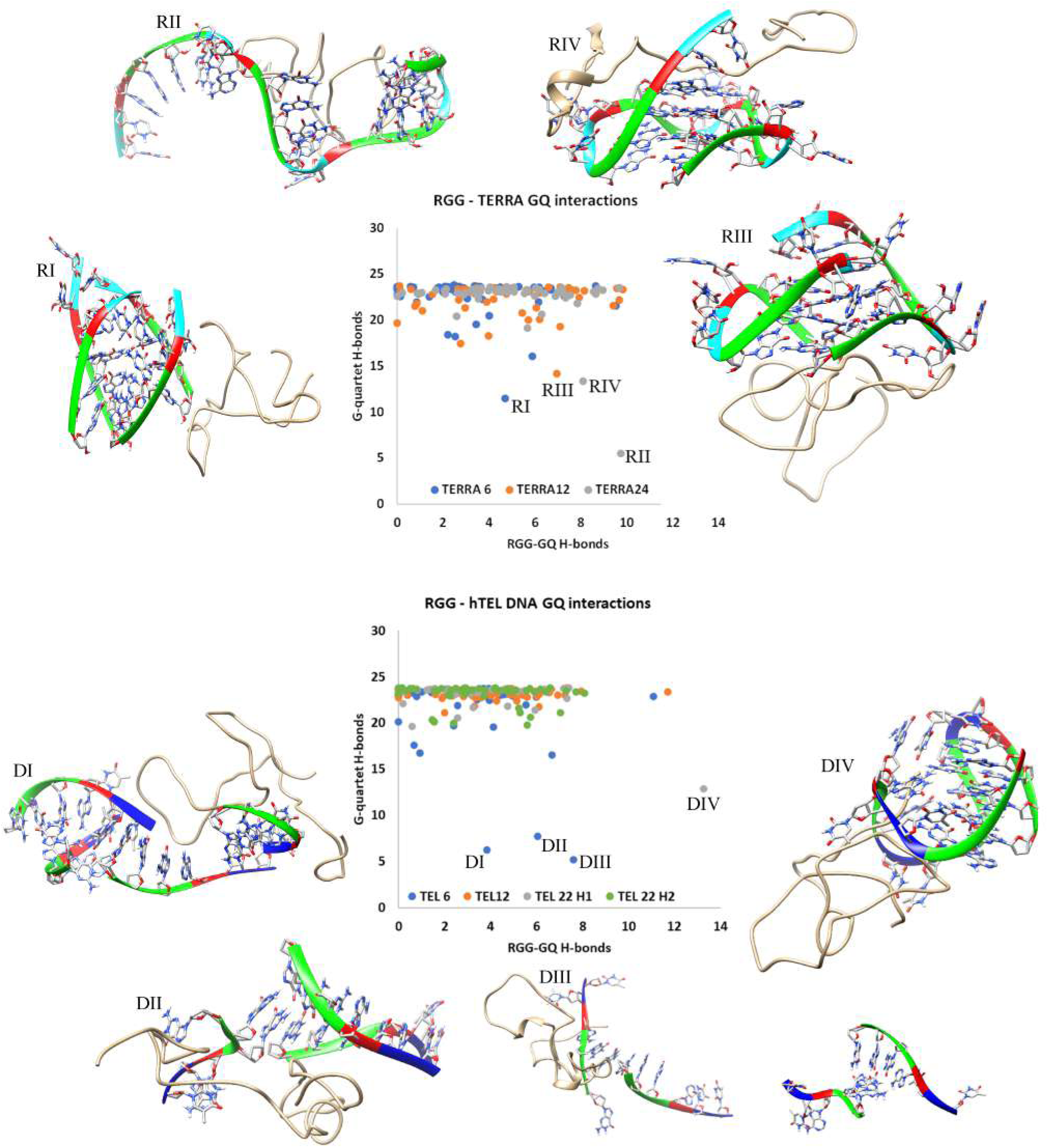
Scatter plot between the number of intermolecular RGG-GQ (RNA and DNA) hydrogen bonds and intramolecular G-quartet hydrogen bonds highlighting the complex stability. The unfolding complexes are labelled and the structures after 50ns MD are shown.

#### 5. Binding mode of RGG defines the mechanism of RGG induced destabilization of GQs

Interesting mechanistic insights into RGG-induced GQ destabilization were observed by analyzing the trajectories of unstable complexes. Instabilities in GQ can be induced by two major mechanisms like, perturbing the core-bound ions or the grooves. The core-bound ions can be disrupted by end stacking interactions, while interactions with the Guanine backbone can disrupt via the grooves. One of the TERRA12 complexes that is unstable with only 15 G-quartet H-bonds, binds RGG in the end stacking mode (labelled RIV in Fig. 7). By visualizing this trajectory, we clearly observed the expulsion of K+ ion by RGG (Fig. 8(a)). A *π*-stacking interaction by Phe222 stabilizes the G12, while the Gly219 is positioned to intrude into the core of GQ, leading to repositioning and expulsion of a K+ ion in the core (Fig. 8(a)).

**FIG. 8:**
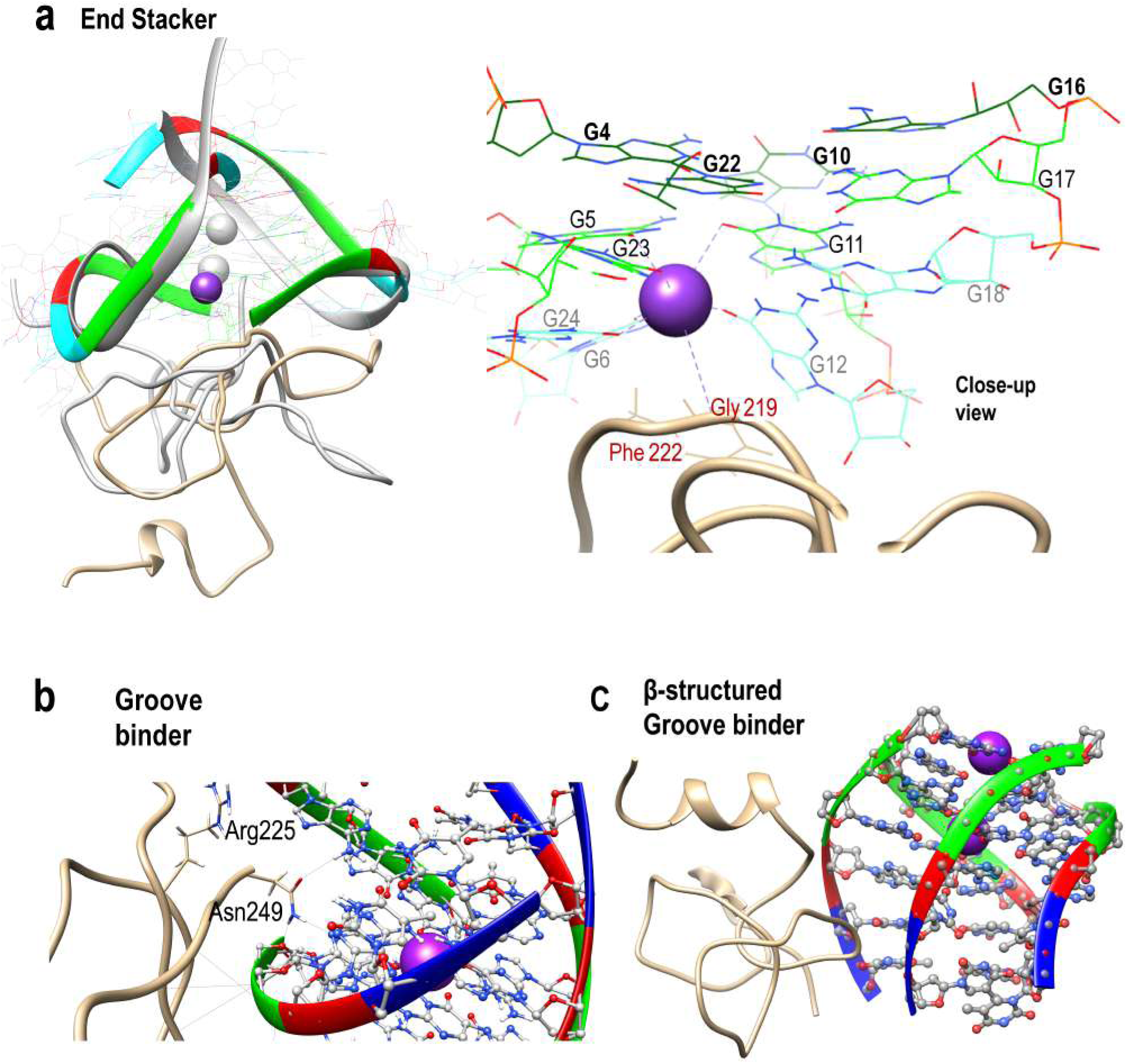
RGG induced unfolding mechanism in (a) the end stacking mode of TERRA12 and (b) groove binding mode of tel6 are shown as examples. The superposition of initial structure (in grey) and 50 ns simulated conformations of TERRA12 are shown to clearly portray the loss of coordinated K+ ions due to RGG. (c) depicts the unfolding of tel6 by a *β*-structured motif binding in the groove. The RGG backbone is shows as ribbons while the sidechains are depicted in wire representation. The K+ ion is shown as a purple sphere and its coordination shell is identified with dotted lines. Phosphate: Yellow, Carbon: gray (RGG)/brown (GQ), Nitrogen: Blue, Oxygen: red, Hydrogen: white and K+ ion: purple.

The tetramolecular structures are particularly easy to unfold by interactions at the grooves. In this mode of binding, the sidechains of RGG amino acids are oriented to interact with the backbone and nucleobases of GQs, straining the GQ to unfold. In one of the tel6 complexes (labelled DIII in Fig. 7), the sidechain of Asn249, interacts directly with an oxygen atom in the deoxyribose sugar of G5 and N3 atom in the Watson-crick face of G4 (Fig. 8(b)). These interactions are due to the protrusion of Asn sidechain into the groove between two parallel strands of DNA, thereby disintegrating the tetramolecular GQ through the grooves. Apart from this, *β*-structured motifs, as shown in Fig.8(c), can bind to the grooves of GQs and induce destabilization. Altogether, our observations indicate that the orientation of RGG sidechains and their ability to form *β*-structured motifs induces desta-bilization in the GQ structure, strengthening the possibility of an RGG-induced mechanism for destabilizing GQs. It has been shown previously that the GQ unfolding rate of individual UP1 domain (*k_obs_* = 0.63 ± 0.01) is lower than UP1+RGG (*k_obs_* = 1.93 ± 0.01)^20^. Our study supports this observation since we show that the individual RGG domain is capable of causing perturbations in the GQ topology, hence catalyzing the unfolding activity of UP1. The observation suggests that the RGG not only acts as an adaptor for recognizing GQ but also plays an augmented role in catalyzing the GQ unfolding.

## III. CONCLUSION

IDRs and IDPs are shown to exhibit a variety of functions by exploiting their inherent disorderliness. They form unique molecular signatures to specifically cater to a definite function. These molecular or structural signatures can be the formation of a particular secondary structure upon binding with a folded partner^33^ or retain the disordered state to recognize the folded partner^34,35,36,37^, and in few cases, express non-specific interactions with another disordered molecule^31^. Such diversity in their structures is necessary to fulfil their diverse functions by recognizing different biomolecules. One such IDR is the RGG-rich domain, which has the ability to bind proteins and nucleic acids. The nucleic acid binding property of RGG-rich regions is well known, and the RGGs of several functionally important IDPs like hnRNPA1, FMRP, and FUS express either stabilizing or destabilizing effects on GQ motifs. Binding studies on FMRP RGG-box with an RNA duplex-quadruplex junction revealed a well-formed *β*-turn in RGG is responsible for its binding to the RNA^38^. A similar involvement of *β*-spiral structure in FUS is reported as necessary for binding a GQ motif^39^. Though the importance of *β*-structures to bind nucleic acids, particularly GQ motifs, is known, a high-resolution mechanistic detail is still lacking. Our study is an attempt to faithfully capture a high-resolution conformational landscape explored by the RGG domain of hnRNPA1 through advanced simulations and further explore its functional significance. Our hybrid replica exchange method has been able to successfully generate the conformational states attained by RGG in solution that also matches well with the chemical shifts of an NMR ensemble. Beyond matching the chemical shifts, our generated conformation ensemble showed secondary structures similar to their propensities in the experimental ensemble. The conformation ensemble generated in our study shows a high propensity for *β*-turns and transient but recurring *β*-sheets. Our docking study highlights that the *β*-turns can be accommodated well in the grooves of GQ.

The RGG of hnRNPA1 is reported to enhance the unfolding of intramolecular human telomeric GQ by UP1 during telomere maintenance. The primary recognition site of RGG is reported as the bases forming the propeller loops of unimolecular GQs. Previous studies have hypothesized that RGG is required for recruiting UP1, however, our study has clearly showed additional purpose for the RGGs. Based on our extensive docking and simulations, we propose an addendum to this hypothesis that the initial binding of RGG might induce local perturbations in GQ, which might facilitate UP1 binding and be responsible for the enhanced unfolding of UP1+RGG. Our study shows that the individual RGGs were able to destabilize GQs, although less frequently (∼8 unfolding events out of the total 350 complexes generated). However, the destabilizing effect of RGG should be attributed to the orientation of sidechains rather than the actual three-dimensional structure. The RGG-GQ interactions are mainly dominated by Phe, Tyr and Asn, and the orientation of these longer sidechains in the GQ grooves is mainly responsible for destabilization. The concentration of hnRNPA1 and GQs, along with physiological conditions, play a vital role in molecular recognition since the RGG conformations and GQ topologies are highly dependent on these factors. Altogether, in this study, we have presented a protocol for generating a high-resolution conformation ensemble of an IDP (the RGG domain of hnRNPA1), capturing the heterogeneity and exploring their functional significance.

## IV. METHODS

### A. Conformer generation of RGG

The initial unfolded structure of RGG domain was obtained from Iterative Threading ASSEmbly Refinement (I-TASSER) which is a widely used server for automated protein structure prediction and structure-based functional annotation^40–42^. Using the amino acid sequence as a starting point, I-TASSER constructs 3D structural models. The generated structure of RGG domain is a collapsed coiled-coil structure. The usage of I-TASSER is limited to generate initial coordinates of the protein and further conformation sampling was carried out using molecular dynamics (MD) simulations.

For IDPs, computational methods can be used to generate a conformation ensemble to study their functions. Some of the commonly used, less computationally demanding methods are flexible-meccano^43^, BUNCH^44^ and IDPConformerGenerator^45^, among others. These methods use backbone dihedrals of non-secondary structural elements compiled from the existing protein structures, folded or disordered. Based on prior knowledge, propensities for secondary structures or long-range interactions can also be included as criteria in these models. However, the most commonly used conformer generation methods are based on molecular dynamics or Monte Carlo simulations. In general, coarse-grained models^46,47^ are used to reduce the time complexity over all-atom models, however, these force fields have several limitations to sample IDPs. We have developed an advanced sampling technique called replica exchange with hybrid tempering (REHT) to generate faithful conformations of IDPs^23^. This method has been used to generate all-atom models and has been tested extensively to generate conformation ensembles of several IDPs like Histatin 5 and *α*-synuclein. We have used this REHT method to generate the conformation ensemble of RGG in this study.

### B. REHT Simulation

Unlike the commonly used temperature based replica exchange, our hybrid approach involves the exchange of both the temperature as well as the Hamiltonian across the replicas. The protein was solvated using the 3-site rigid TIP3P water model in a cubic box with a minimum distance of 1 nm from the surface of the protein. The systems were also neutralized to maintain a physiological concentration of NaCl (0.15 M). The Charmm36m^48^ forcefield was used to model the protein. The simulations were carried out using Gromacs-2016.5 patched with Plumed-2.4.1^49–51^. Initially, the modelled protein was energy minimized using steepest descent algorithm for 50,000 steps to avoid any poor contacts. The energy minimized structure was then thermalized and equilibrated sequentially in NVT and NPT ensembles for 1 ns.

Next, the protein and the solvent were coupled separately to the target temperatures using the Nose-Hoover thermostat and the final production simulation was performed in the NVT ensemble. A cut-off of 1 nm was used for calculating the electrostatic and vdw interactions and Particle Mesh Ewald was used for long-range electrostatics. To integrate the equations of motions, the Leap-frog integrator with the time step of 2 fs was used. LINCS algorithm was used to constrain all the hydrogen atoms. Exchanges were attempted at every 1 ps interval. In this approach, the Hamiltonian of the solute particle in the different replica is scaled-down to 0.6-1 while also raising the temperature of the systems simultaneously up to a maximum of 500 K. This way, as a function of both lambda scaling and explicit thermostat conditions, a very high temperature is ensured to be realized on the protein solute to sufficiently overcome the energy barriers. The solvent is heated up mildly in order for the rapid reorientation of the hydration shell upon conformational change of the protein. 20 replicas were used to simulate the RGG in a temperature range of 300 K to 500 K for a cumulative period of 1 *µ*s and an acceptance probability of 0.3. Post processing analyses of the trajectories were performed with the Gromacs analysis tools and the trajectories were visualized using VMD.

### C. Chemical shift calculation

We have used SPARTA+(version 2.90) in conjunction with a software package called Mdtraj^52,53^. SPARTA+ employs a well-trained neural network algorithm to construct quantitative relations between chemical shifts and protein structures, including backbone and side-chain conformation, H-bonding, electric fields and ring-current effects. It’s single-level feed-forward multilayer artificial neural network (ANN) is capable of identifying the dependence of ^15^*N ^H^*, ^1^3*CO*, ^13^*C^α^*, ^13^*C^β^*, ^1^*H^α^*, ^1^*HN* chemical shifts on the local structural and dynamic information as well as amino acid type, and those of its immediate neighbors. Md-traj is a python software package which is designed to analyse trajectory data generated from MD simulations. We have compared our predicted data with the experimentally calculated chemical shift submitted in Biological Magnetic Resonance Data Bank (BMRB entry 50017)^26,54^. For comparison purposes, we have used secondary chemical shift data by removing random coil chemical shifts from the raw chemical shift. For ΔC*α*-ΔC*β* calculation, the secondary chemical shifts of ^13^*C^α^* and ^13^*C^β^* are subtracted in order to do away with the referencing error, if there is any.

### D. Secondary structure propensity calculation

We have used the secondary structure prediction algorithm (SSP) developed by Prof. Julie D. Forman-Kay’s group**^?^**. We have used ^13^*Cα*, ^13^*Cβ* and 1*^H^α* chemical shifts for this purpose as recommended by the algorithm developers in the context of intrinsically disordered proteins. For random coil and secondary structure chemical shifts and standard deviations, we have used RefDB. All the data have been re-referenced.

### E. t-SNE clustering

We have chosen a nonlinear dimensionality reduction method called t-distributed Stochastic Neighbor Embedding (t-SNE) in order to separate high dimensional heterogeneous IDP ensemble into meaningful clusters. An important aspect of t-SNE algorithm is the perplexity value, a tuneable parameter in t-SNE that balances the information between the local and global features of the dataset under investigation. Diligent choice of perplexity value for a given data is important to be able to most discretely divide the data into unambiguous clusters. In our work, we have noticed that for very low perplexity values, the clustering is extremely diffusive in nature and for a high value of perplexity parameter, the entire dataset seems to be treated as a single cluster. The performance of t-SNE is fairly robust to changes in the perplexity. In this work, we have explored several perplexity value options ranging from 50-2000. We have chosen the values which most discretely divides the datapoints into separate clusters. To aid the result of t-SNE clustering in further partitioning, we have employed a widely used centroid based technique called K-means clustering which is one of the favoured and standard unsupervised machine learning algorithms. We have used SciKit tool for these analyses which is an open source library for machine learning in Python^55^.

### F. Structures of RGG and GQ

The 50 conformations of RGG obtained by clustering the REHT trajectories were docked with the human telomeric DNA and RNA GQ (also known as GQ). Unimolecular, bimolecular and tetramolecular forms of both DNA and RNA quadruplexes were modeled in this study (Table I). The tetramolecular TERRA (or TERRA6) was modeled using the solution structure with PDB ID: 6GE1, while the unimolecular TERRA24 was modeled based on the telomeric DNA quadruplex with PDB ID: 1KF1. The NMR structure of bimolecular telomeric RNA (or TERRA12) is available in the Protein Databank with the ID 2KBP. The structures of all forms of telomeric DNA quadruplexes were taken directly from the Protein Databank with IDs 1NB9 (tetramolecular or tel6), 1K8P (bimolecular tel12), 2HY9 (unimolecular form 1 tel22h1) and 2JPZ (unimolecular form 2 tel22h2). The unimolecular telomeric DNA in K+ ion solution adopts two different hybrid conformations that are equally populated and both the structures (differentiated as tel22h1 and tel22h2) were taken for our docking studies. The numbers behind TERRA* or tel* indicates the length of each RNA or DNA strand forming the GQ. Additional bases were added to the 5’ or 3’ ends of the GQ structures to match the sequences used in ITC studies by Ghosh et al^20,22^. Two K+ ions were placed manually between the three G-quartets of all 7 GQs. The modeled structures were simulated for a period of 300 ns and the structures extracted after 10 ns simulation were used as starting structures for docking studies.

### G. HADDOCK Docking protocol

HADDOCK^56^ utilizes a data-driven approach for molecular docking. The stand-alone version of HADDOCK 2.4 was used to drive the docking of RGG with telomeric RNA and DNA GQs. Since the RGG conformations are randomly coiled structures of 53 amino acids, all residues are highly solvent exposed. In order to perform an unbiased docking study, the distance between the center of mass of the protein and GQ alone were used as a restraint criterion. The docking process is performed in three steps: rigid-body docking, semi-flexible refinement and water refinement. About 10,000 complexes were generated during the first stage rigid-body docking by using the standard HADDOCK protocol. Among these, 400 lowest-energy structures were selected for subsequent semi-flexible simulated annealing and explicit solvent (water) refinement, to optimize the interface. HADDOCK scoring was performed according to the weighted sum (HADDOCK score) of different energy terms, including the van der Waals energy, electrostatic energy, distance restraints energy, inter-vector projection angle restraints energy, diffusion anisotropy energy, dihedral angle restraints energy, symmetry restraints energy, binding energy, desolvation energy and buried surface area. The final structures were clustered using the fraction of common contacts (FCC) with a cutoff of 0.6 and a minimum cluster size of 4.

### H. Simulation of RGG-GQ complexes

For interaction studies of RGG-GQ complexes (listed in Table I), we performed molecular dynamics simulation using Gromacs-2020.4^57^. We used a99SBdisp forcefield of Paul Robestelli for RGG, OL15 force field for DNA quadruplexes and OL3 force field for RNA quadruplexes. The RGG-GQ complex systems (obtained from HADDOCK) were solvated with TIP3P water in a periodic box extending up to 1 nm and 1.2 nm in all directions respectively. The systems were neutralized and additional ions were added to mimic a salt concentration of 0.15 M. The short-range interactions were truncated with a cut-off distance of 1 nm. Electrostatic interactions were treated by particle-mesh Ewald with a real space cut-off value of 1 nm. Bonds containing hydrogens were constrained using the LINCS algorithm. The solvated and neutralized systems were energy minimized using the Steepest Descent algorithm followed by an equilibration of 5 ns and subsequently production runs. The temperature and pressure of the systems were maintained at 310 K and 1 atm using the nose-hoover thermostat and Parrinello-Rahman barostat in an NPT ensemble. Post processing analyses were performed with the Gromacs analysis tools, CPPTRAJ module of AmberTools20 and the trajectories were visualized using Visual Molecular Dynamics^58^, UCSF Chimera v1.13. RGG-GQ interaction matrices were consolidated by calculating all heavy-atom contacts within 0.4 nm that are present for atleast 1% of the total simulation period for each GQ (50 complexes * 50 ns = 2.5 *µ*s).

## V. DATA AVAILABILITY

Input files needed to initiate molecular simulations and full trajectory data of all simulations for all systems considered in this work are available on our server for download. The server data can be publicly accessed via our Figshare repository here.

## AUTHOR CONTRIBUTIONS

AS conceived the idea. AS and RA designed and formulated the project with help of SB and IR. IR constructed and set up the hnRNPA1-LCD systems and SB constructed and set up models for hnRNPA1-GQ complex systems. IR and SB carried out all simulations and generated all the data, figures and plots for the manuscripts. SB, IR, RA and AS analyzed the data. IR and SB created the first draft of the manuscript, AS refined the draft with help of IR, SB and RA.

## ACKNOWLEDGEMENTS

IR would like to acknowledge financial support from Ministry of Education, Government of India. R.A. would like to thank the early career fellowship from the DBT-Wellcome Trust India Alliance (Grant number: IA/E/18/1/504308). S.B. would like to thank the National Supercomputing Mission grant for financial support. AS acknowledges the financial support from the Indian Institute of Science-Bangalore and the high-performance computing (HPC) facility ”Beagle” setup from grants by a partnership between the Department of Biotechnology of India and the Indian Institute of Science (IISc-DBT partnership programme). AS also thanks the DST for the National Supercomputing Mission grants (DST/NSM/R&D-HPC-Applications/2021/03.10 and DST/NSM/R&D-HPC-Applications/Extension Grant/ 2023/27) for the HPC support. FIST program sponsored by the Department of Science and Technology and UGC, Centre for Advanced Studies and Ministry of Human Resource Development, India. AS would also like to thank the Teams Science Grant from the DBT-Wellcome Trust India Alliance (Grant number: IA/TSG/21/1/600245) and the DBT National Network Project (NNP) grant (BT/PR40323/BTIS/137/78/2023 grants. This work was initiated through the Matrics grants (MTR/2023/001040) from the Science and Engi-neering Board (SERB), India and both authors thanks SERB for this support.

## Supporting Information

**Fig. S 1:**
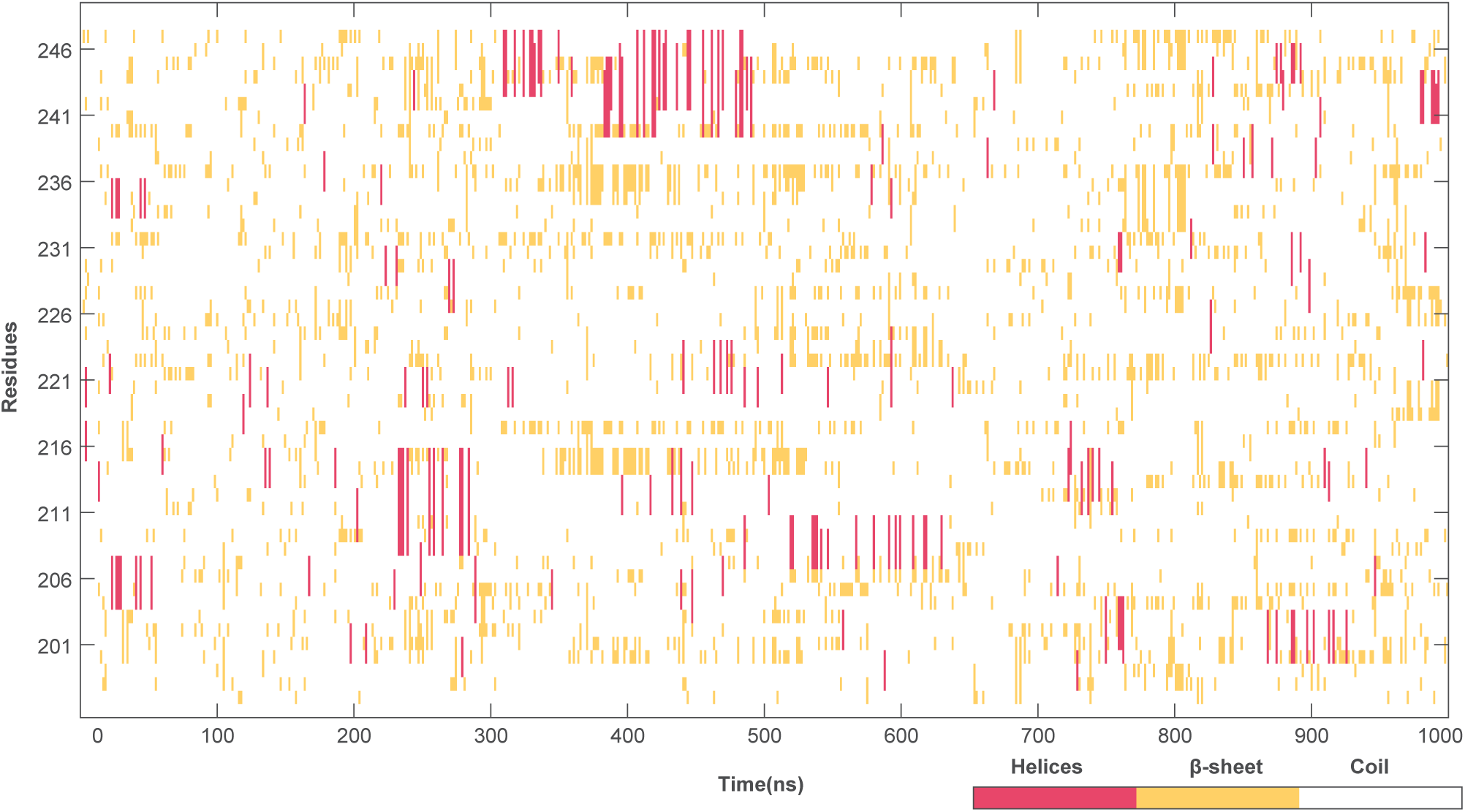
Time evolution of secondary structures formed during simulation calculated using the DSSP algorithm. The time (1000ns) is plotted on X-axis. Y axis contains the residues. Relevant colour codes for the secondary structures are: White-Coil, Yellow-*β*-sheet, Red-Helices.

**Fig. S 2:**
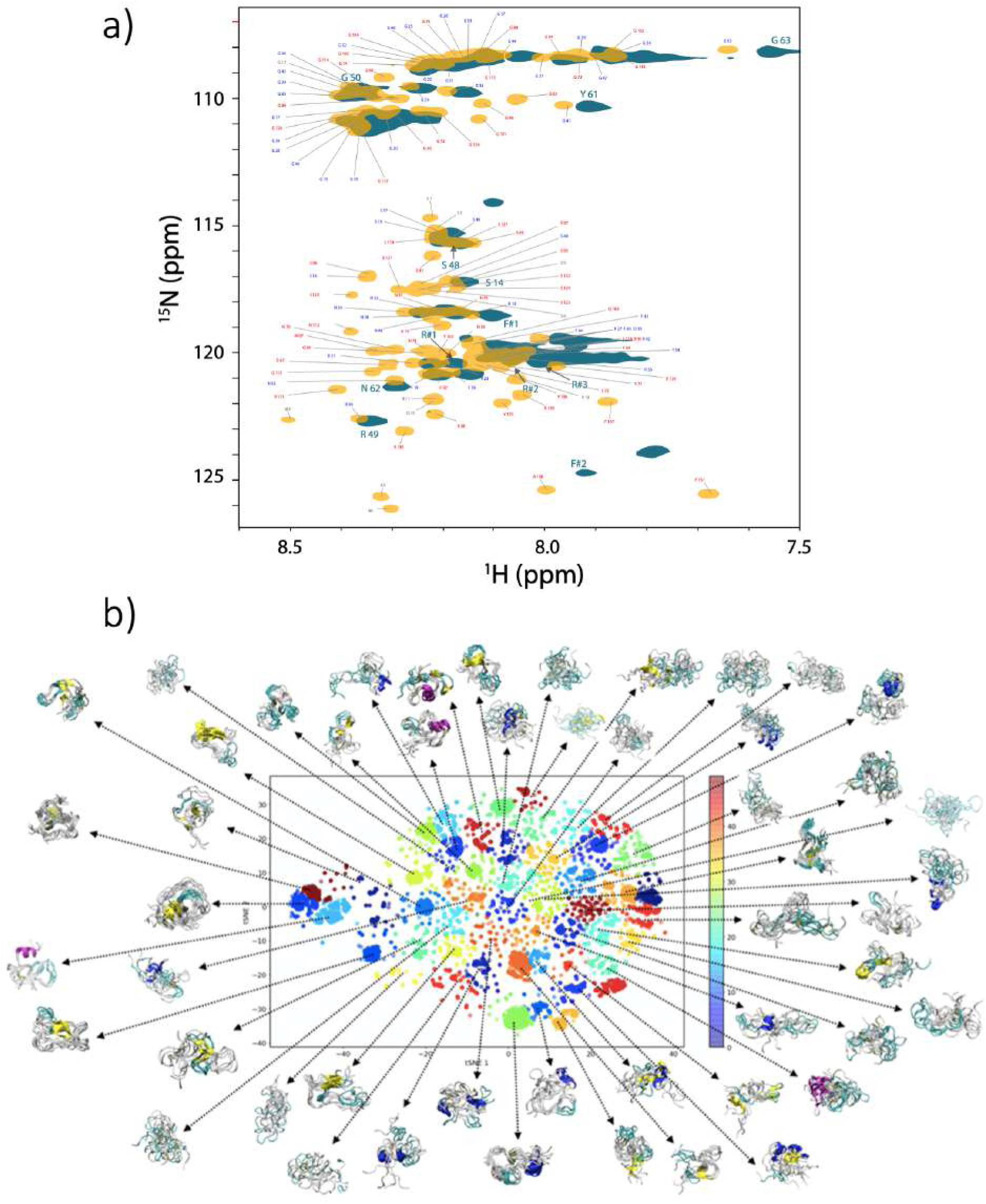
a) Merged ^15^*N* −^1^ *H* HQSC plot. Yellow markers represents full hnRNPA1-LCD^1^. Blue colored labels indicate RGG domain, red colored labels indicate PrLD domain. Teal markers represent only RGG domain represents.^2^ b)t-SNE clustering of RGG-box domain.The 50 clusters are color coded and representative structures from each clusters and superposed and mapped to the corresponding clusters.

**Fig. S 3:**
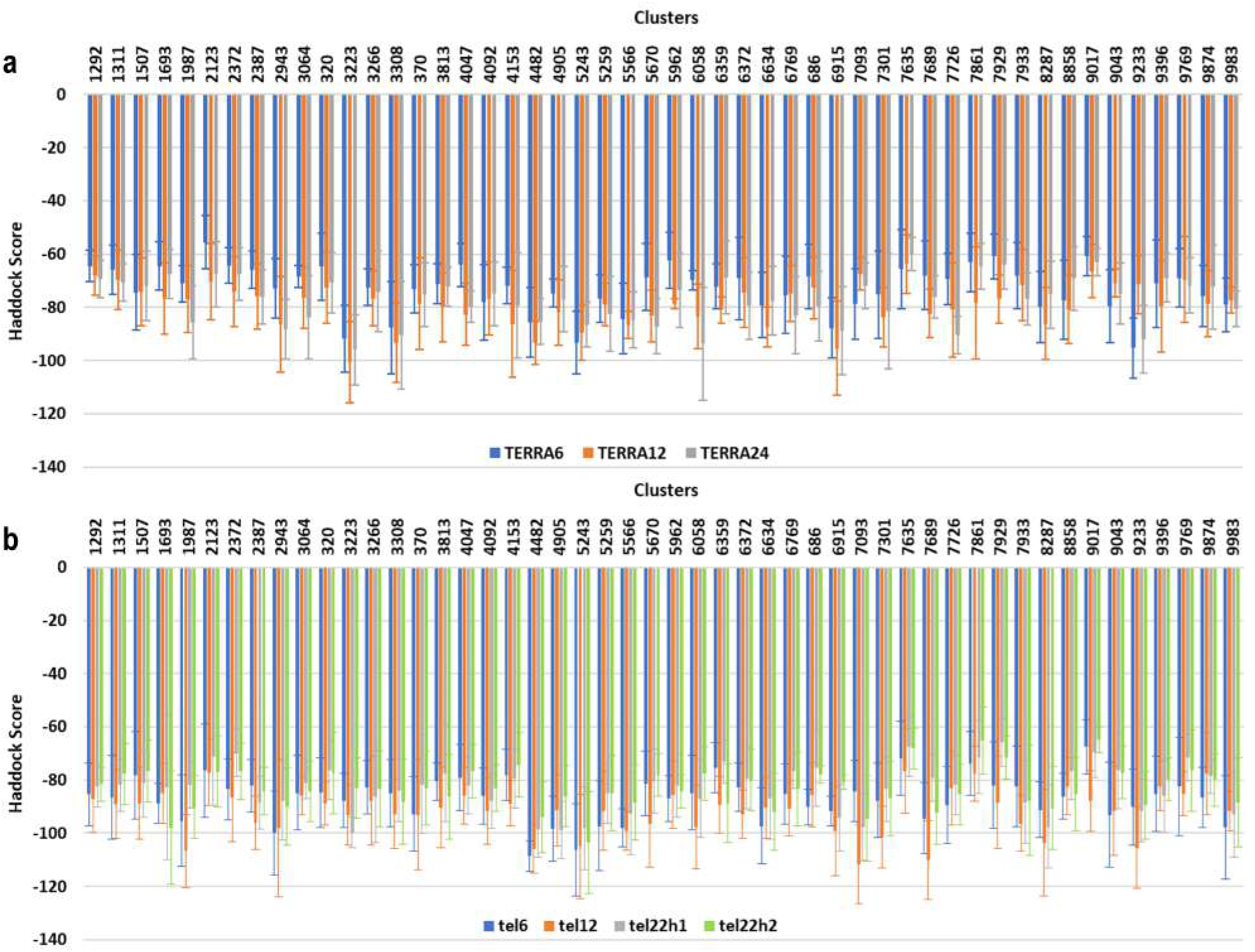
The Haddock scores for each of the 50 RGG conformers used in the docking study with the seven telomeric G-quadruplexes (GQ), (a) tetramolecular (TERRA6 in Blue), bimolecular (TERRA12 in Orange) and unimolecular (TERRA24 in Gray) RNA quadruplexes. (b) tetramolecular (tel6 in Blue), bimolecular (tel12 in Orange), unimolecular form 1 (tel22h1 in Gray) and unimolecular form 2 (tel22h2 in Green) DNA quadruplexes.

**Fig. S 4:**
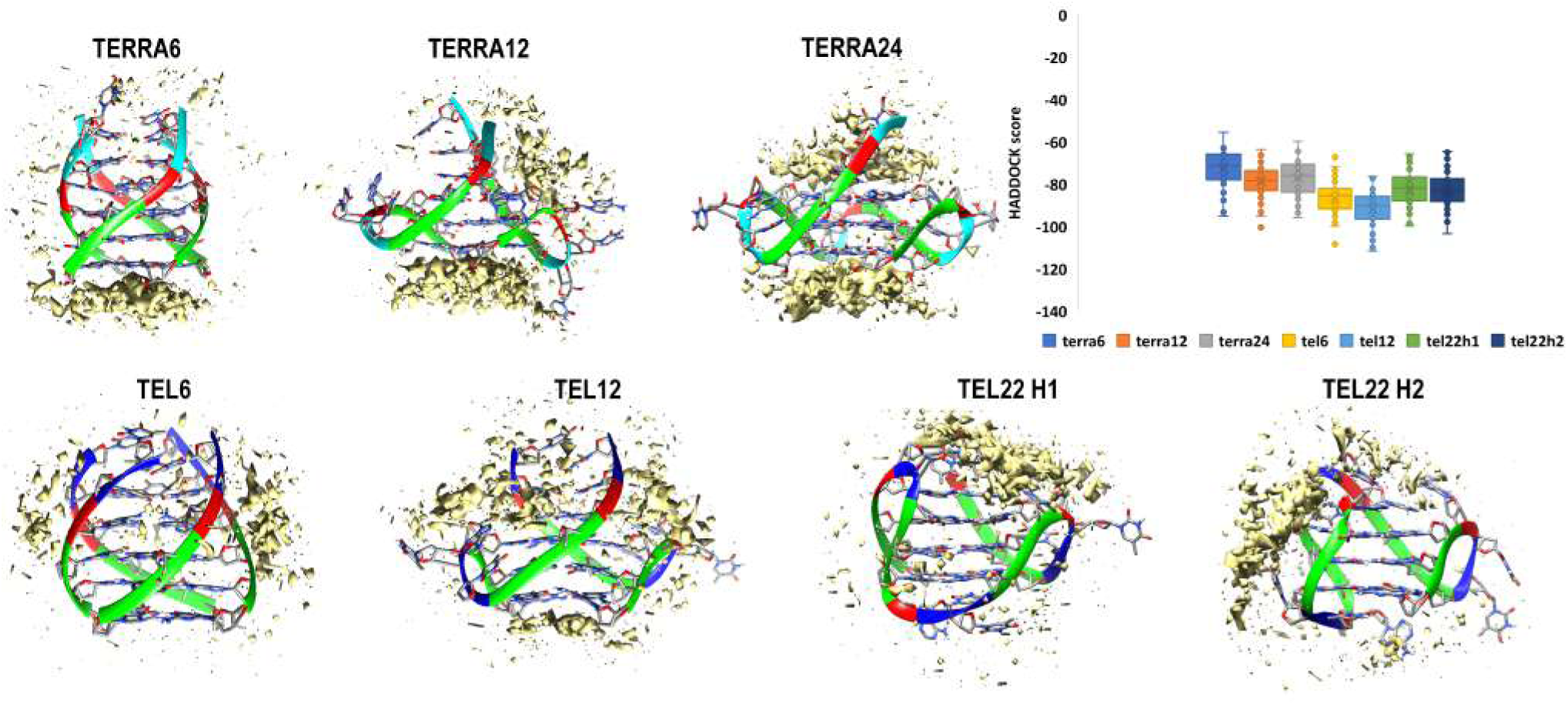
Occupancy map depicting the extent of RGG-GQ interaction and the high density of interactions at specific sites on GQ are shown as an isosurface mimicking the first solvation shell (Yellow surface). The range of HADDOCK scores obtained for the docking of RGG with the seven GQ structures. Boxplots indicate minimum, median, maximum, and upper and lower quartiles. The GQ backbone is shown as ribbons while the bases are displayed as sticks. The bases are colored as Green: Guanine, Red: Adenine, Blue: Thymine and Cyan: Uracil.

**Fig. S 5:**
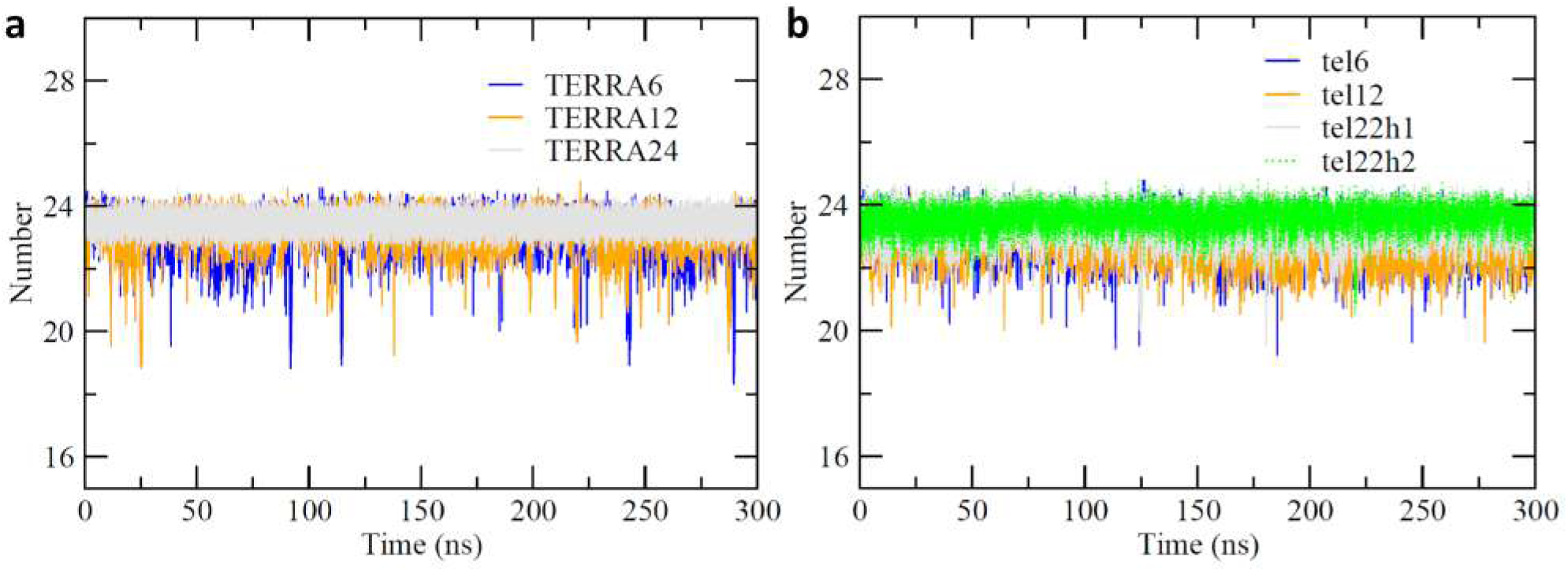
Stability of free RNA (a) and DNA (b) GQ monitored by calculating the number of hydrogen bonds within the three G-quartets.

## SUPPLEMENTARY TABLES

**Table S I:**
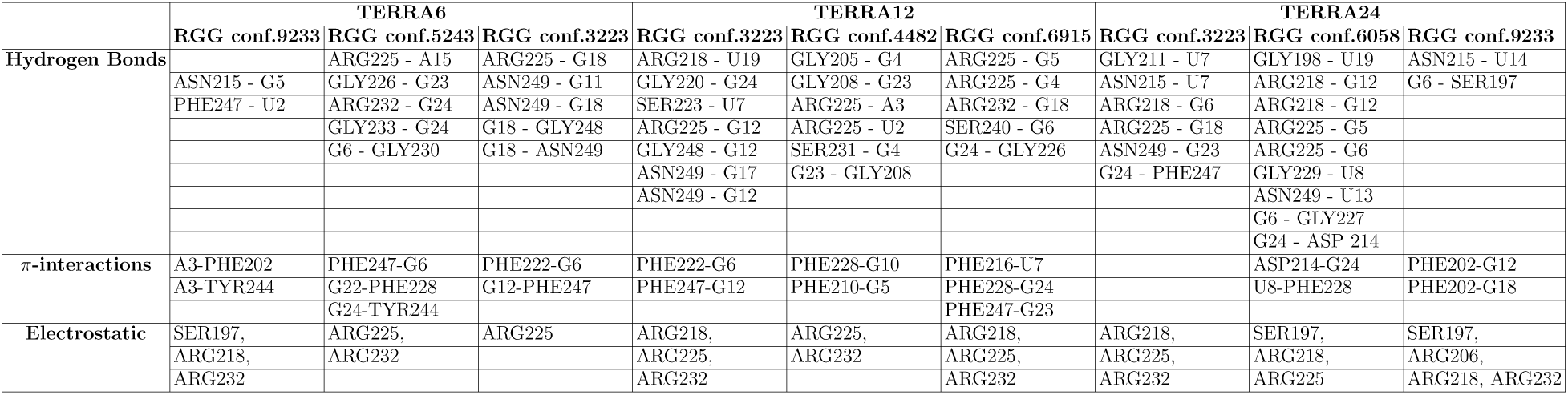
RGG-RNA GQ interactions identified in the best three complexes screened by the HADDOCK score.

**Table S II:**
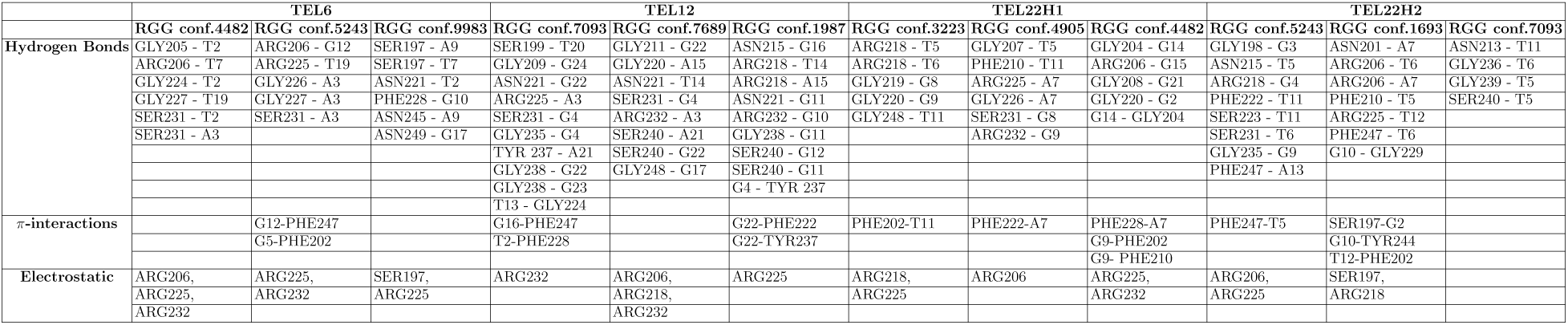
RGG-DNA GQ interactions identified in the best three complexes screened by the HADDOCK score.

